# Screening and Identification of Lujo Virus Entry Inhibitors from an FDA-Approved Drugs Library

**DOI:** 10.1101/2021.09.01.458657

**Authors:** Junyuan Cao, Yang Liu, Siqi Dong, Minmin Zhou, Jiao Guo, Xiaoying Jia, Yueli Zhang, Yuxiao Hou, Gengfu Xiao, Wei Wang

**Author notes:** Address correspondence to Wei Wang. Junyuan Cao, Yang Liu, and Siqi Dong contributed equally to this work.

## Abstract

The Lujo virus (LUJV) belongs to the Old World (OW) genus *Mammarenavirus* (family Arenaviridae); it is categorized as a biosafety level (BSL) 4 agent. Currently, there are no U.S. Food and Drug Administration (FDA)-approved drugs or vaccines specifically for LUJV or other pathogenic OW mammarenaviruses. Here, a high-throughput screening of an FDA-approved drug library was conducted using pseudotype viruses bearing LUJV envelope glycoprotein (GPC) to identify inhibitors of LUJV entry. Three hit compounds, trametinib, manidipine, and lercanidipine, were identified as LUJV entry inhibitors in the micromolar range. Mechanistic studies revealed that trametinib inhibited LUJV GPC-mediated membrane fusion by targeting C410 (located in the transmembrane (TM) domain), while manidipine and lercanidipine inhibited LUJV entry by acting as calcium channel blockers. Meanwhile, all three hits extended their antiviral spectra to the entry of other pathogenic mammarenaviruses. Furthermore, all three could inhibit the authentic prototype mammarenavirus, lymphocytic choriomeningitis virus (LCMV), and could prevent infection at the micromolar level. This study shows that trametinib, manidipine, and lercanidipine are candidates for LUJV therapy, and highlights the critical role of calcium in LUJV infection. The presented findings reinforce the notion that the key residue(s) located in the TM domain of GPC provide an entry-targeted platform for designing mammarenavirus inhibitors.

**IMPORTANCE:** To date, only one LUJV outbreak has been recorded; it occurred in 2008 and resulted in a fatality rate of 80% (4/5 cases). Pathogenesis studies and therapeutic strategies are therefore urgently needed. Repurposing approved drugs can accelerate the development of drug design and facilitate the understanding of infectious mechanisms. Here, three compounds, trametinib, manidipine, and lercanidipine, were identified as entry inhibitors against LUJV. Studying the underling mechanisms revealed that a key residue (C410) in LUJV GPC modulates its sensitivity/resistance to trametinib and demonstrated the critical role of calcium in LUJV infection.

## INTRODUCTION

The Lujo virus (LUJV) was identified in South Africa in 2008; it is the pathogen of Lujo hemorrhagic fever (LUHF) (1). LUJV is an enveloped, negative-sense, bi-segmented RNA virus belonging to the genus *Mammarenavirus* (family Arenaviridae) (1, 2). There are 39 unique *Mammarenavirus* species currently recognized by the International Committee on Taxonomy of Viruses. The original classification of mammarenaviruses, which was based mainly on virus genetics, serology, antigenic properties, and geographical relationships, divided them into new world (NW) and old world (OW) mammarenaviruses (3). Based on phylogenetic analysis, LUJV has been demonstrated to be distinct from OW and NW mammarenaviruses and has been reported to use neuropilin-2 (NRP2) as its primary receptor, rather than the typical receptors utilized by OW (α-dystroglycan) and NW (transferrin receptor) mammarenaviruses (4, 5). LUJV; the OW Lassa virus (LASV); and some NW mammarenaviruses, including the Junín virus (JUNV), Machupo virus (MACV), Guanarito virus (GTOV), Chapare virus (CHAPV), and Sabiá virus (SBAV), are known to cause severe hemorrhagic fever; they are listed as biosafety level (BSL) 4 agents (6, 7).

The mammarenavirus RNA genome encodes viral polymerase, nucleoproteins, matrix protein (Z), and glycoprotein complex (GPC). GPC is synthesized as a polypeptide precursor that is sequentially cleaved by signal peptidase and the cellular protease subtilisin kexin isozyme-1/site-1 protease to generate the three subunits of the mature complex: the retained stable signal peptide (SSP), the receptor-binding subunit GP1, and the membrane fusion subunit GP2 (8–11). These three non-covalently bound subunits form a (SSP/GP1/GP2)_3_ trimeric complex that is present at the surface of the mature virion; it plays an essential role in virus entry. The relatively conserved mammarenavirus SSP and GP2 form an interface that not only contributes to the stabilization of the prefusion conformation of GPC, but also provides an “Achilles’ heel” that can be targeted by the entry inhibitors.

To date, no vaccines or specific antiviral agents against LUJV have been developed. To address this issue, here we screened a U.S. Food and Drug Administration (FDA)-approved drug library of 1,775 compounds. Drug repurposing is a strategy that is used to accelerate the discovery and development of new and emerging pathogens. Meanwhile, understanding the antiviral mechanisms of potential drugs that could combat LUJV could provide novel insights into its pathogenesis. The safety, pharmacokinetics, and mechanisms of approved drugs have been intensively investigated. Here, therefore, we screened drugs that targeted the entry step of LUJV infection, as this could block viral replication and spreading at an early stage. After three rounds of screening, trametinib, manidipine, and lercanidipine were found to be highly effective against LJUV entry; thus, they could offer potential new therapies to treat LUHF.

## RESULTS

### High-throughput screening (HTS) for LUJV entry inhibitors

As studies of LUJV require BSL-4 equipment, we utilized a replication-competent recombinant virus of LUJV (LUJVrv, comprising a vesicular stomatitis virus (VSV) backbone with a genome containing green fluorescent protein and LUJV GPC) for HTS. LUJVrv possesses the entry characteristics of LUJV and can be handled under BSL-2 conditions (12, 13). The HTS assay conditions, including the seeding cell density and LUJVrv infective dose, were optimized at 1.5 × 10^4^ cells and 1.5 × 10^2^ PFU per 96-well plate, respectively. Under these optimized conditions, the coefficient of variation (CV) and *Z*’ factor were 4.21% and 0.95, respectively, demonstrating that this assay represents a promising approach for the large-scale screening of inhibitors. A schematic of the HTS is shown in Fig. 1A. Inhibitors were defined as prime candidates with >90% inhibition and no apparent cytotoxicity at a concentration of 10 μM. Of the 1,775 tested compounds, 145 (8.17%) were considered as prime candidates. Then, the prime candidates were subjected to counter-screening to rule out the inhibition of VSV genome replication. Twenty-one compounds (1.18%) were selected with inhibition of VSV < 10% at 25 μM. Screening was then performed to reconfirm the results using these compounds over a broader concentration range (0.024–75.0 μM). Four compounds (0.23%) were selected based on their concentration-dependent inhibitory effects. Among these four compounds, the top three hits (trametinib, manidipine, and lercanidipine) were selected for further investigation; ospemifene was eliminated because of its cytotoxicity (Fig. 1B).

**Fig. 1.**
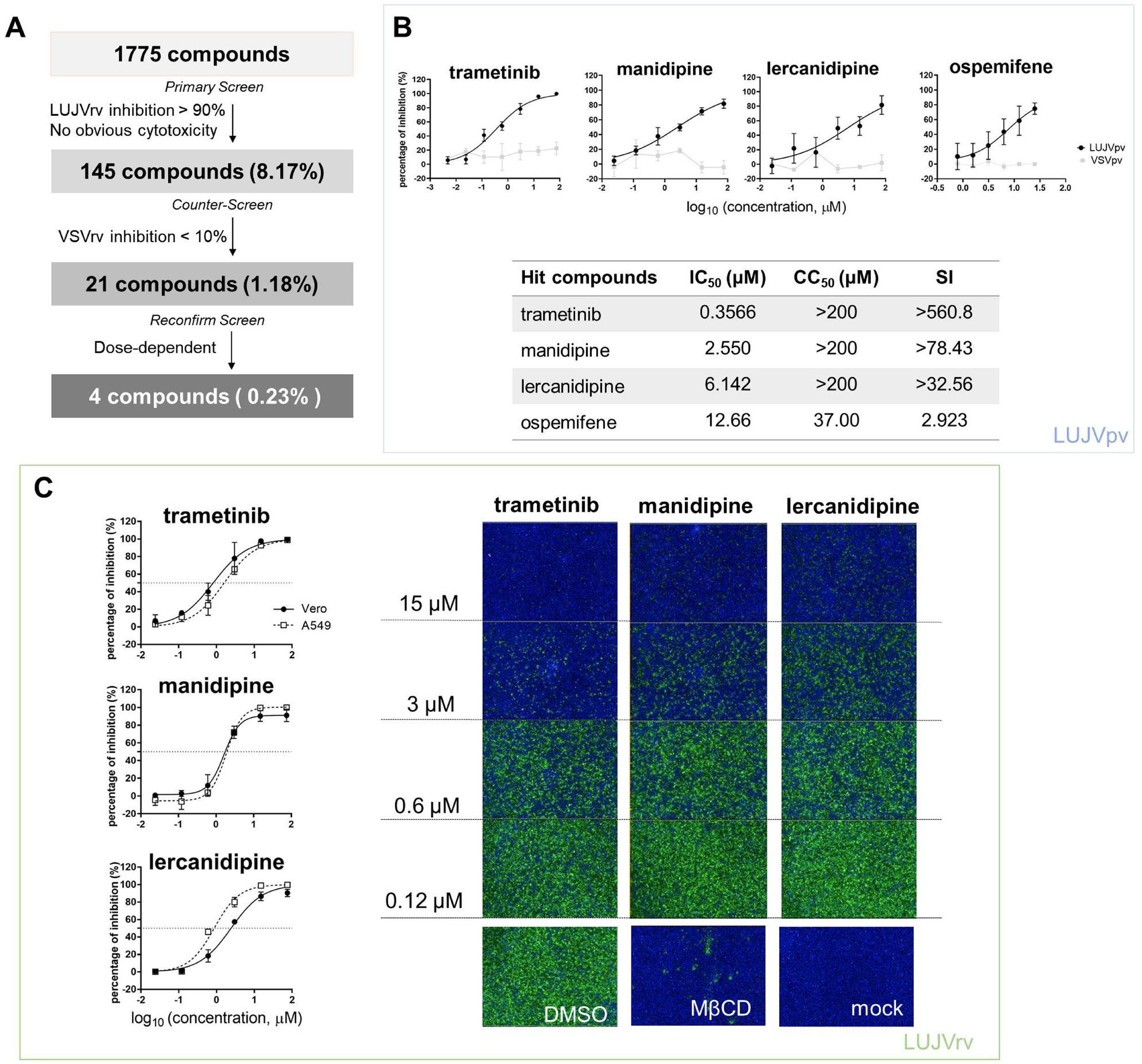
HTS for inhibitors of LUJV entry from an FDA-approved drug library. (A) HTS assay flowchart. (B) (Top) Dose-response curves of four hit compounds against LUJVpv. Cells were seeded at a density of 1.5 × 10^4^ cells per well in 96-well plates. After overnight incubation, cells were treated in duplicate with each compound at the indicated concentrations; 1 h later, cells were infected with LUJVpv (MOI, 0.01) for 1 h. The infected cells were lysed 23 h later, and their luciferase activities were measured. (Bottom) IC_50_, CC_50_, and SI values of the hit compounds. Cell viability was evaluated using MTT assay. (C) (Left) Dose-response curves of trametinib, manidipine, and lercanidipine against LUJVrv on both Vero and A549 cells. (Right) Images showing the inhibition of the top three hits against LUJVrv infection. Vero cells were preincubated with compounds at 37 °C for 1 h, followed by incubation with LUJVrv (MOI, 0.1) in the presence or absence of compounds for 1 h. GFP-positive cells were counted using an Operetta high-content imaging system 23 h later. Data are presented as means ± SD, from five to eight independent experiments.

Trametinib is an inhibitor of mitogen-activated protein kinase (MAPK), while manidipine and lercanidipine are dihydropyridine (DHP) voltage-gated Ca^2+^ channel antagonists. We evaluated the 50% inhibitory concentration (IC_50_) of the hit compounds using LUJVpv (Fig. 1B). Furthermore, the inhibitory effects were confirmed using LUJVrv on both Vero and A549 human epithelial cell lines (Fig. 1C). To validate the antiviral effects, trametinib, manidipine, and lercanidipine were purchased from other commercial sources and tested; the cytotoxic and antiviral effects were similar to the results shown in Fig. 1.

### Trametinib inhibits LUJV GPC-mediated membrane fusion

The unique retained SSP, together with GP2, provides an “Achilles’s heel” that can be targeted by many arenavirus entry inhibitors. We carried out a membrane fusion assay to test whether the three hit compounds act by targeting the SSP-GP2 interface. Notably, in addition to transfecting 293T cells with LUJV GPC, plasmids expressing pEGFP and CD63_GY234AA_ (a mutant leading to CD63 localization on the cell surface) were also transfected. This indicates the formation of syncytium and the triggering of LUJV GPC-mediated membrane fusion (4, 5). As shown in Fig. 2A, low pH induced obvious membrane fusion in cells transfected with both GPC and mutated CD63, whereas no syncytium formed in cells treated with neutral pH, or in cells lacking CD63_GY234AA_.

**Fig. 2.**
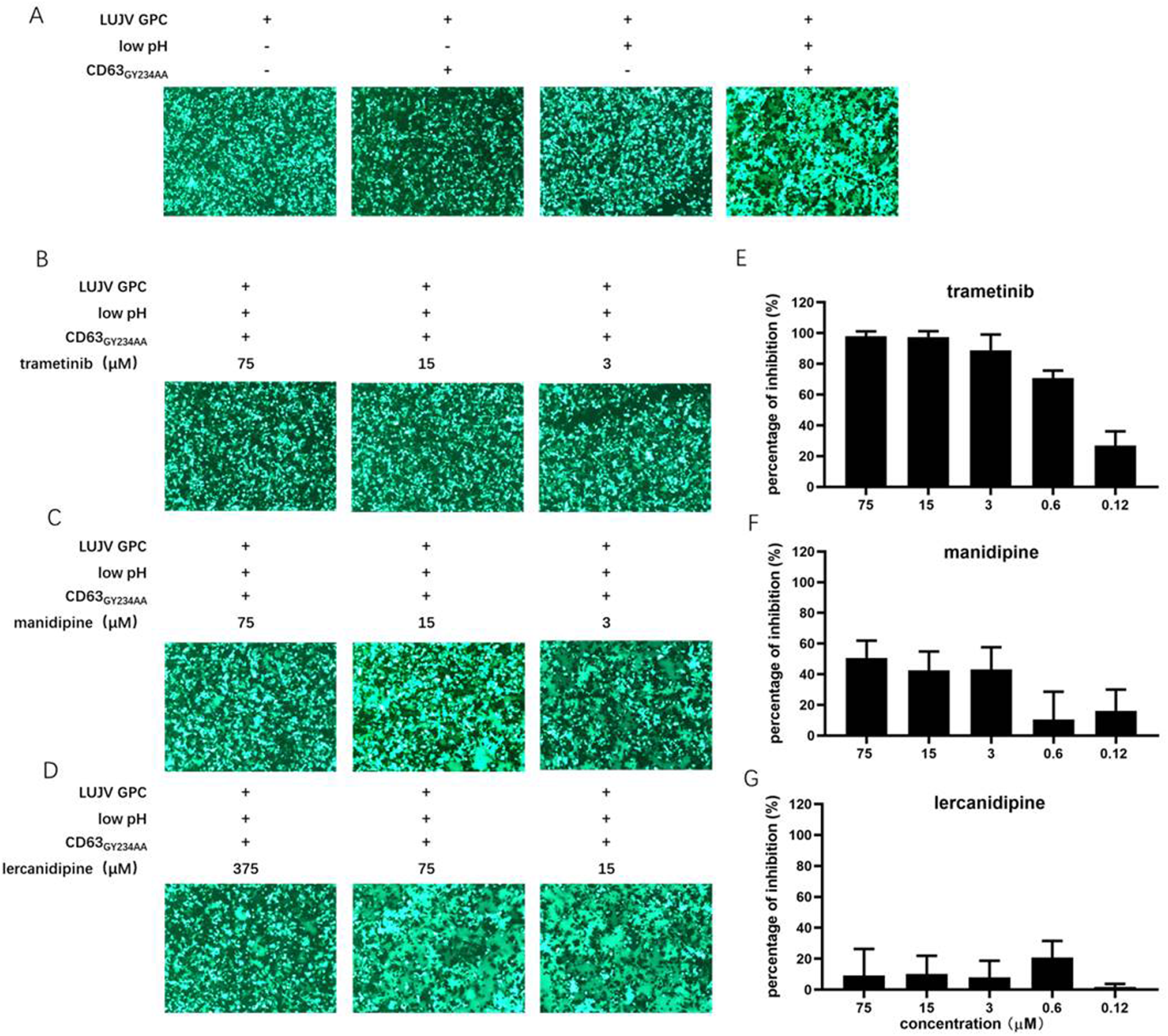
Inhibition of LUJV GPC-mediated membrane fusion with trametinib. (A) LUJV GPC-mediated membrane fusion was triggered by the low pH in the presence of membrane-anchored CD63. 293T cells co-transfected with pCAGGS-LUJV GPC, pcDNA 3.1-CD63_GY234AA_, and pEGFP-N1 were treated with acidified (pH 5.0) medium for 20 min. The cells were then washed and incubated with neutral medium, and syncytium formation was imaged 1 h later. (B-D) Qualitative evaluations of the effects of trametinib (B), manidipine (C), and lercanidipine (D) on LUJV GPC-mediated membrane fusion. Images are representative fields from three independent experiments. (E-G) Quantitative analysis of the effects of trametinib (E), manidipine (F), and lercanidipine (G) on LUJV GPC-mediated membrane fusion using ImageJ. Data are presented as means ±SD from three or four independent experiments.

As shown in Fig. 2B and C, trametinib significantly inhibited syncytium formation under all tested concentrations (3 to 75 μM). Mild inhibition was observed under treatment with manidipine even at the highest tested concentration (75 μM). Notably, lercanidipine exerted little inhibition on membrane fusion, even at higher concentrations (15–375 μM; Fig. 2D). Although the three hit compounds had similar IC_50_ values (all < 10 μM), they might inhibit LUJV entry through different mechanisms. Trametinib blocks LUJV entry by inhibiting GPC-mediated membrane fusion, but the other two calcium channel antagonists might use alternative mechanism(s).

A luciferase-based fusion assay has previously been used to quantify membrane fusogenicity (13–16). However, this luciferase-based fusion assay failed to evaluate LUJV GPC-mediated membrane fusion, as there no difference in luciferase activity was observed between the acidic pH treatment group and the control group. One possible reason for this might be that one additional plasmid, (CD63_GY234AA_) needed to be co-transfected with other plasmids (13–16), which led to a low transfection efficacy. To this end, here we used ImageJ to quantify the average size of each syncytium. In line with the qualification analysis, trametinib exerted dose-dependent inhibition on the LUJV GPC-mediated membrane fusion (Fig. 2B and E); neither manidipine nor lercanidipine inhibited this effect (Fig. 2C, D, F, and G).

### Mutation C410A confers resistance to trametinib

As most arenavirus entry inhibitors target the SSP-GP2 interface to prevent membrane fusion, we further investigated the viral targets of the hit compounds. We selected the adaptive mutant virus by serially passaging the replication, competing with LASVrv, in the presence of the hit compounds. The starting concentration was 10 μM for trametinib and manidipine and 20 μM for lercanidipine, corresponding to the IC_90_ values obtained in the LUJVrv inhibition assay (Fig. 1C). Parallel passaging of LUJVrv was also conducted in dimethyl sulfoxide (DMSO) as a control. In the trametinib-treated group, resistance was detected after three rounds of passaging; it increased in passage P4 and then remained stable through passages P5 to P6 (Fig. 3A). To identify the viral genetic determinant(s) responsible for the resistance, we sequenced the trametinib-resistant virus and identified the cysteine (C)-to-glycine (G) mutation at amino acid position 410, which is located at the C terminus of the transmembrane domain (TM) on GP2 (Fig. 3B). To confirm that the C410G mutation conferred trametinib resistance, we introduced a mutation into the GPC gene and generated a pseudotype mutant virus. Compared with the wild type (WT) LUJVpv, the C410G mutant virus exhibited robust resistance to trametinib, but remained sensitive to manidipine and lercanidipine (Fig. 3C). Resistance was confirmed in the membrane fusion assay by the obvious formation of syncytium in the GPC_C410G_ group, when treated with trametinib. Meanwhile, syncytium formation was robustly inhibited in WT (Fig. 3D).

**Fig. 3.**
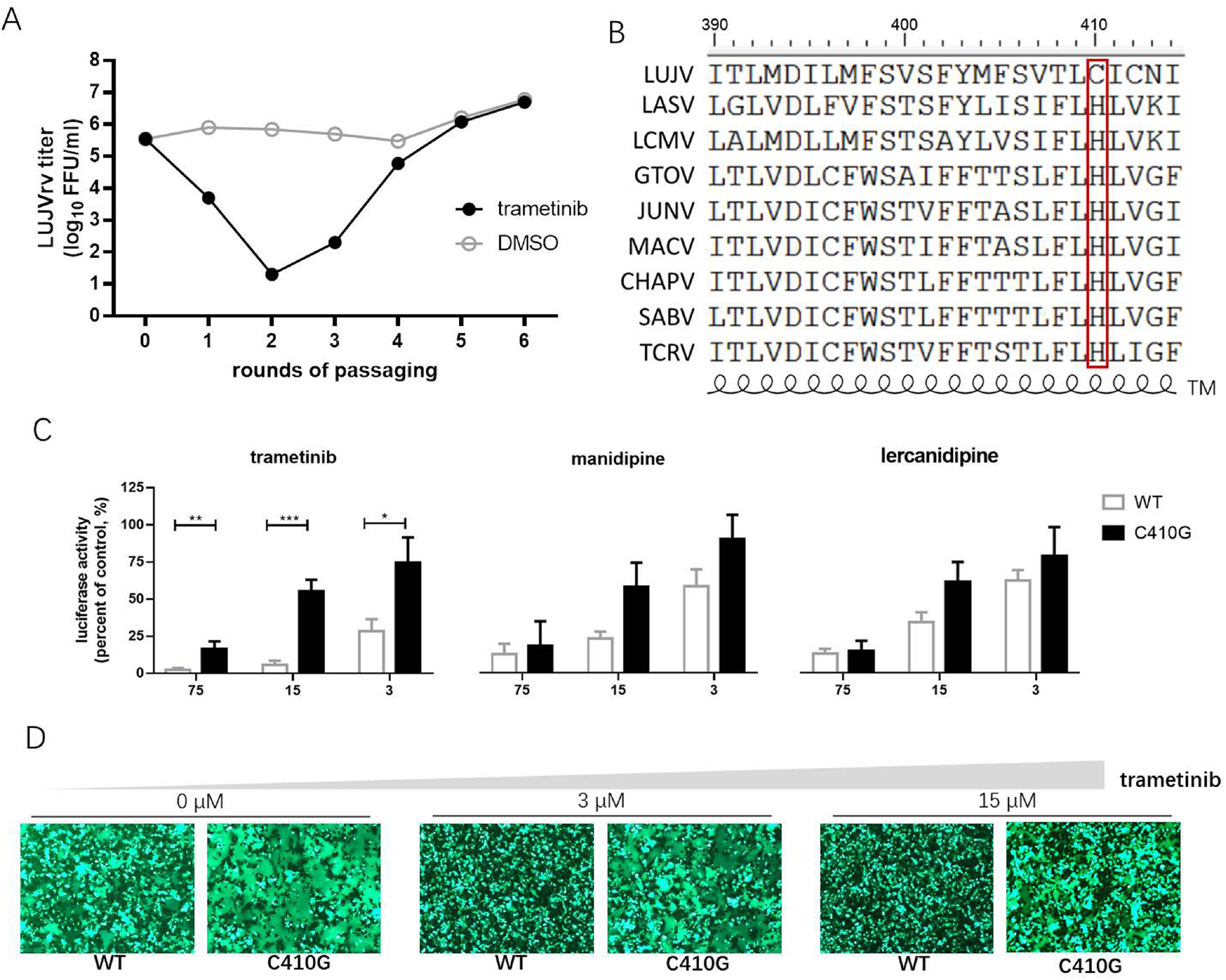
Selection of trametinib-resistant LUJVrv. (A) The adaptive mutant virus was selected by serially passaging LUJVrv in the presence of 10 μM trametinib. In a parallel experiment, LUJVrv passaging in vehicle served as a control. After six rounds of passaging, no further improvement in resistance was detected, and the selection was terminated. (B) Amino acid sequence alignment of the mammarenavirus TM. (C) Resistant activities to trametinib of the WT and C410G LUJVpv. Vero cells were treated with compounds at the indicated concentrations; 1 h later, WT or C410G LUJVpv (MOI, 0.1) was added to the culture for 1 h. The infected cells were lysed 23 h later and the luciferase activities were measured. Data are presented as means ± SD, from two or three independent experiments (***, *P* <0.001; **, *P* <0.01; and *, *P* <0.05). (D) C410G conferred resistance to the inhibition of trametinib against LUJV GPC-mediated membrane fusion. Images are representative fields from three independent experiments.

The cysteine residues located within the border region between the transmembrane domain and the cytoplasmic tail of glycoprotein were usually found to be palmitoylated (17). Furthermore, LUJV GPC possessed the motif C_410_IC, which was reported to be palmitoylated in Ebola virus (EBOV) and Marburg virus (MARV) glycoprotein (18). Thus, we hypothesize that both C410 and C412 might serve as potential palmitoylated sites in LUJV GPC, and that palmitic acid might be the target of trametinib. To this end, C412 was substituted with G, and its drug sensitivity/resistance was evaluated. As shown in Fig. 4A, both the single C412G mutant and the combined C410G/C412G mutant still underwent membrane fusion with acid triggering, similar to that of C410G (Fig. 3D). When treated with trametinib, fusogenicity was inhibited in the C412G group, but not in the C410G/C412G group, suggesting that only C410 served as the viral target of trametinib. Furthermore, LUJV_C412G_pv was still as sensitive to trametinib as that of WT, whereas LUJV_C410G/C412G_pv was resistant to trametinib, similar to LUJV_C410G_pv (Fig. 4B). This confirms that only C410G contributed to the resistance to trametinib.

**Fig. 4.**
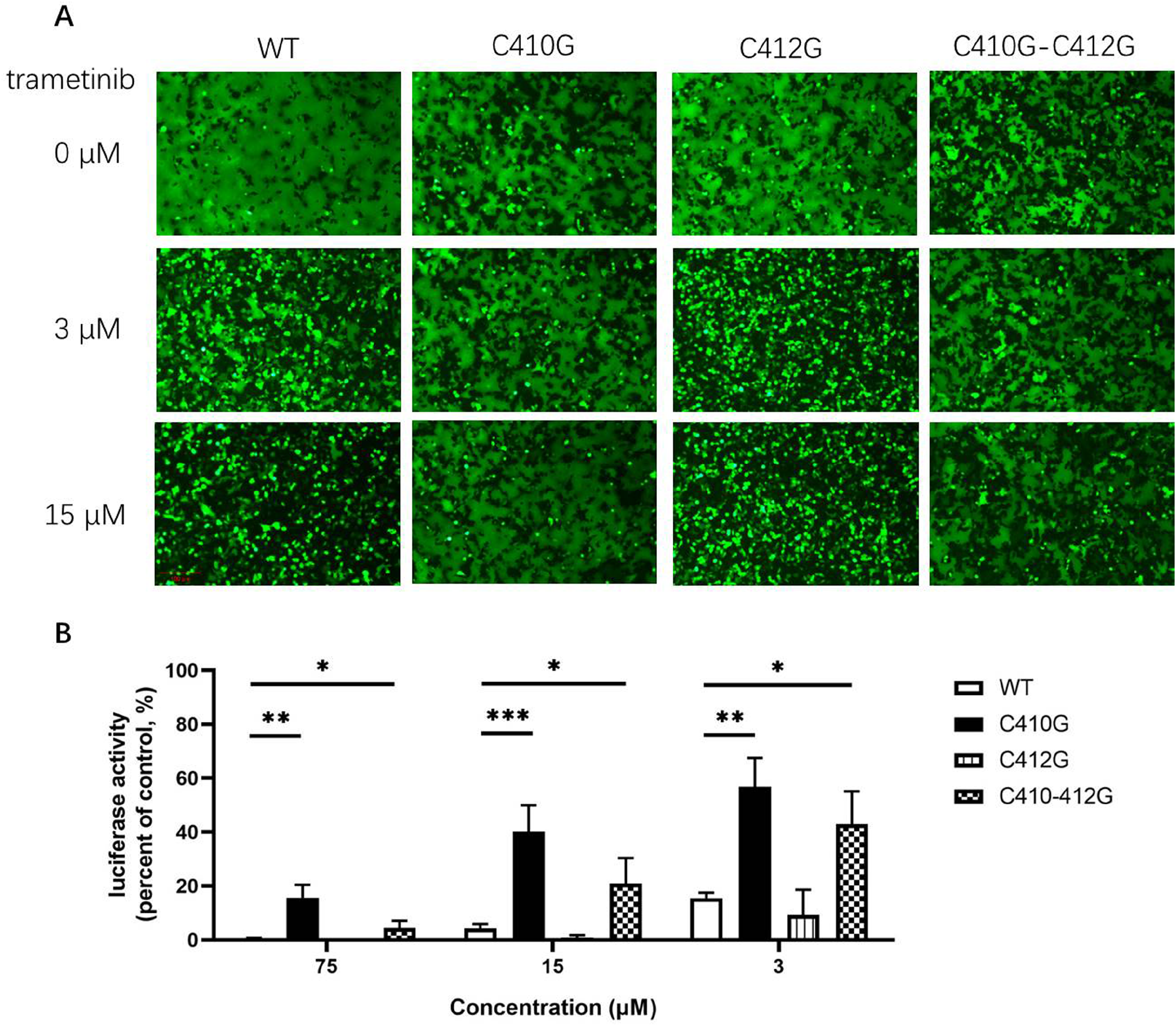
Only C410G, but not C412G, conferred resistance to trametinib. (A) Both C410G and C410G/C412G mutants conferred resistance to the inhibition of trametinib, against LUJV GPC-mediated membrane fusion. Images are representative fields from three independent experiments. (B) Both C410G and C410G/C412G mutants conferred the resistance to trametinib, while both WT and C412G retained their sensitivity. Data are means ± SD from three to four independent experiments.

Notably, we attempted to identify the resistance of LUJVrv to manidipine and lercanidipine. However, after 12 rounds of passaging with increasing concentrations of either compound (10-50 μM for manidipine, 20-50 μM for lercanidipine), the drug-resistant phenotype was absent in both groups, and no adaptive mutation was obtained for either compound. Combined with the finding that manidipine and lercanidipine had little effect on GPC-mediated membrane fusion, we thus speculate that manidipine and lercanidipine inhibited virus entry by targeting cellular compounds, rather than viral proteins.

### Effects of hit compounds against other mammarenaviruses

We further tested whether the hit compounds exhibited their inhibitory effects against the entry of other pseudotypes of other pathogenic mammarenaviruses, such as LASV, lymphocytic choriomeningitis virus (LCMV), Mopeia virus (MOPV), JUNV, MACV, GTOV, SBAV, and CHAPV. Intriguingly, although C410 was found to be unique among the mammarenaviruses (Fig. 3B), trametinib inhibited all of the tested pseudotypes of mammarenaviruses, with IC_50_ values ranging from 0.756 to 10.36 μM (Fig. 5A–5H). We used the BSL-2 level-compatible LCMV Cl-13 strain to test the antiviral effects of trametinib against an authentic mammarenavirus. As shown in Fig. 5I, trametinib inhibited the authentic LCMV infection, with an IC_50_ value of 3.919 μM, similar to that of LCMVpv (4.822 μM, Fig. 5H).

**Fig. 5.**
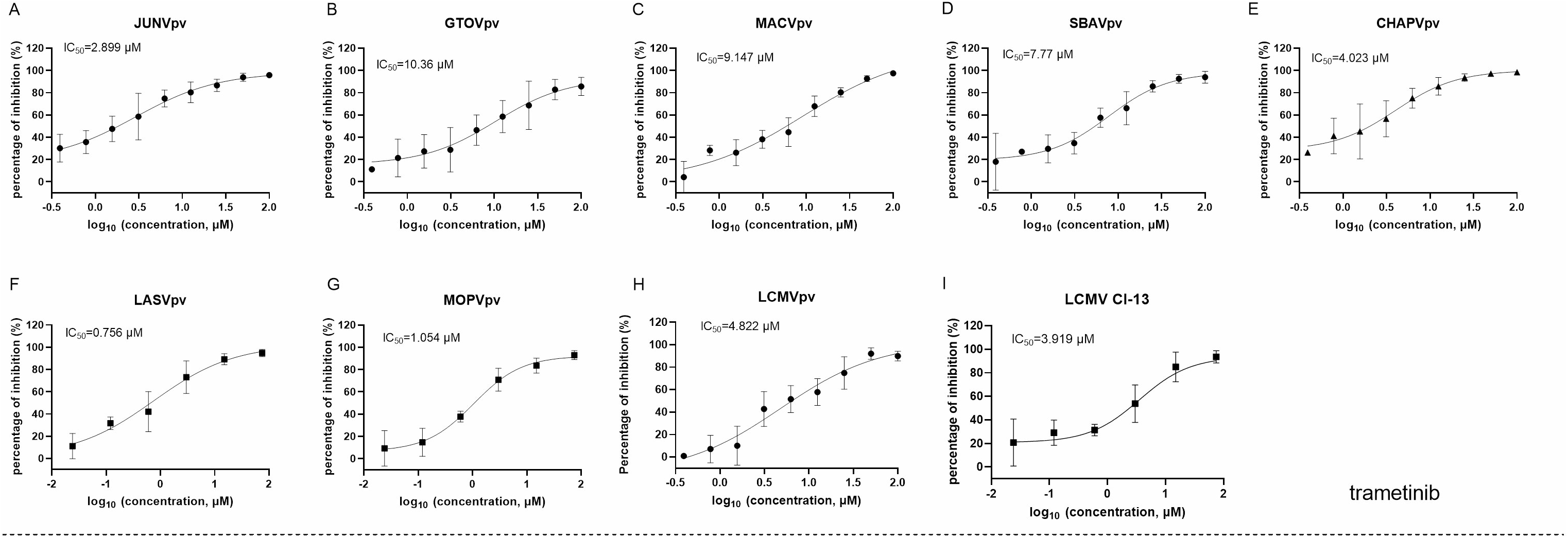
Broad-spectrum antiviral activity of trametinib. (A-H) The inhibition of trametinib against the infection of the pseudotype of mammarenaviruses. (I) Inhibition of trametinib against authentic LCMV infection. Vero cells were incubated with trametinib for 1 h, and LCMV-Cl 13 (MOI, 0.01) was added into the cells 1 h. Then, the supernatants were replaced with medium containing trametinib. After 24 h, the cell lysates were subjected to real time (RT)-qPCR. (J-Q) Effects of manidipine on the infection of the pseudotypes of mammarenaviruses. (R) Inhibition of manidipine against authentic LCMV infection. (S-Z) Effects of lercanidipine on the infection of the pseudotypes of mammarenaviruses. (I) Inhibition of lercanidipine against authentic LCMV infection. Data are presented as means ± SD from three to five independent experiments.

Trametinib has been reported to repress the expression of transferrin 1 (19), which might result in the inhibition of NW mammarenavirus entry (Fig. 5A–5E). For the other OW mammarenaviruses that do not possess the corresponding C410, we investigated the viral target by serially passaging LASVrv in the presence of 10 μM trametinib, corresponding to the IC_85_ value obtained in the LASVpv inhibition assay (Fig. 5F). As shown in Fig. 6A, resistance emerged after two rounds of passaging. Sequencing of trametinib-resistant LASVrv revealed that the viral target was F446L, which is located in the TM of GP2 (Fig. 6B). F446 is highly conserved in mammals, and has been reported to confer resistance to other structurally distinct membrane fusion inhibitors (15). The sensitivity/resistance was further tested in the presence of trametinib by introducing the F446L mutant into LASVpv, MOPVpv, and LCMVpv. As shown in Fig. 6C to E, LASV_F446L_pv, MOPVpv, and LCMVpv (containing the corresponding F446L mutant) conferred resistance to trametinib, confirming that F446 served as the viral target of trametinib in LASV, MOPV, and LCMV.

**Fig. 6.**
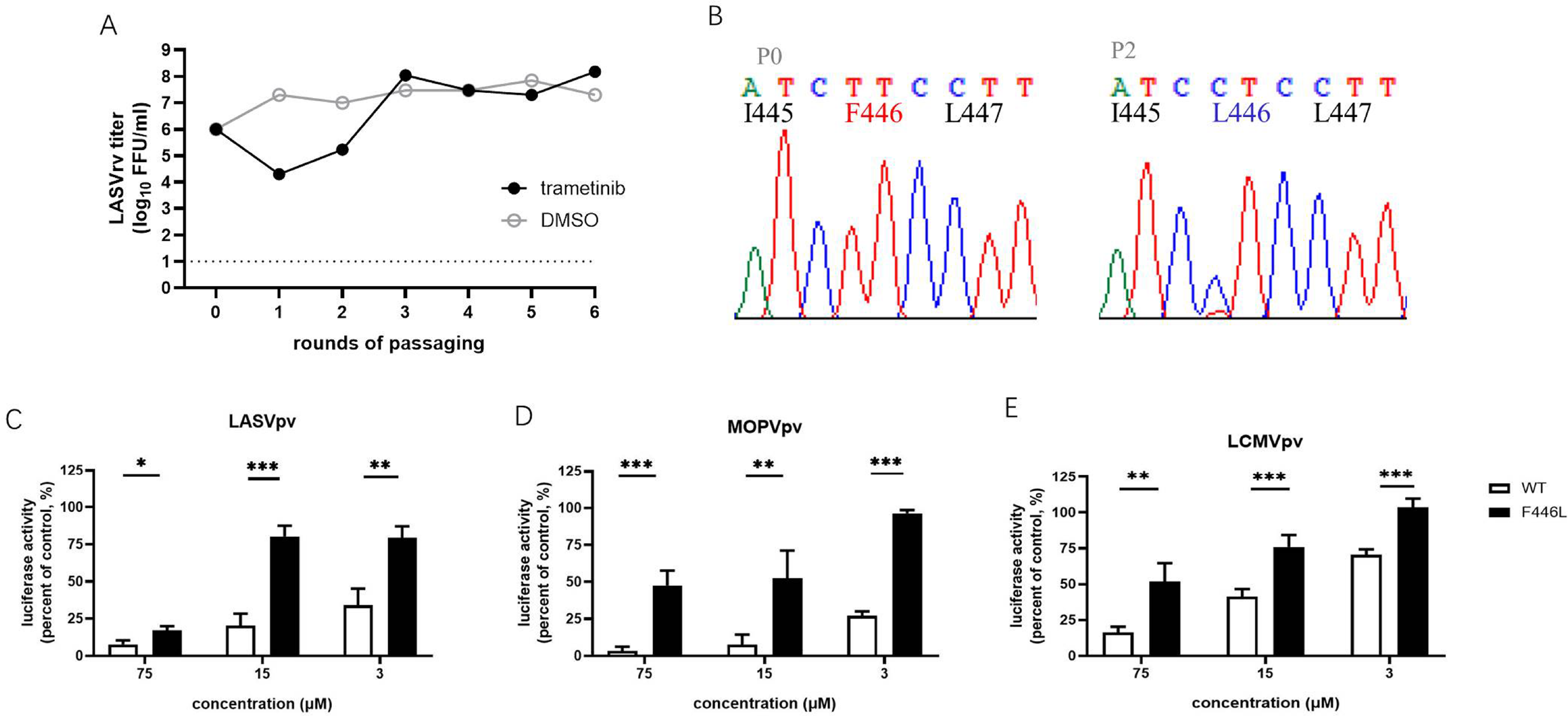
F446L mutant in transmembrane domain of OW mammarenavirus conferred resistance to trametinib. (A) The adaptive mutant virus was selected by serially passaging LASVrv in the presence of 10 μM trametinib. In a parallel experiment, the passaging of LUJVrv in vehicle served as a control. (B) Sequencing chromatograms of P0 and P2 viruses with the highlighted adaptive mutant. (C) LASV_F446L_pv conferred resistance to trametinib. (D) The corresponding F446L mutant MOPVpv conferred resistance to trametinib. (E) The corresponding F446L mutant LCMVpv was still sensitive to trametinib. Data are presented as means ± SD from three to four independent experiments (***, *P* <0.001; **, *P* <0.01; *, *P* <0.05).

We also investigated the broad-spectrum antiviral effects of manidipine and lercanidipine. Both showed robust inhibition against NW mammarenavirus entry, with IC_50_ values ranging from ~ 0.1-1 μM. At the tested concentration, the dose-response curves of both manidipine and lercanidipine for NW viruses showed the typical “s” model, with 2-log spans (Fig. 7A–E, Fig. 7J–N). These results are in line with published reports, in which voltage-gated calcium channel (VGCC) antagonists were found to be critical for the cellular binding and entry of the NW arenaviruses JUNV and Tacaribe virus (20, 21). Notably, although manidipine and lercanidipine exerted ambiguous inhibition against OW pseudo-viruses (Fig 7F–H, Fig 7O–Q), both showed promising inhibition against authentic LCMV infections, with IC_50_ values of 2.344 μM for manidipine (Fig. 7I) and 1.093 μM for lercanidipine (Fig. 7R).

**Fig. 7.**
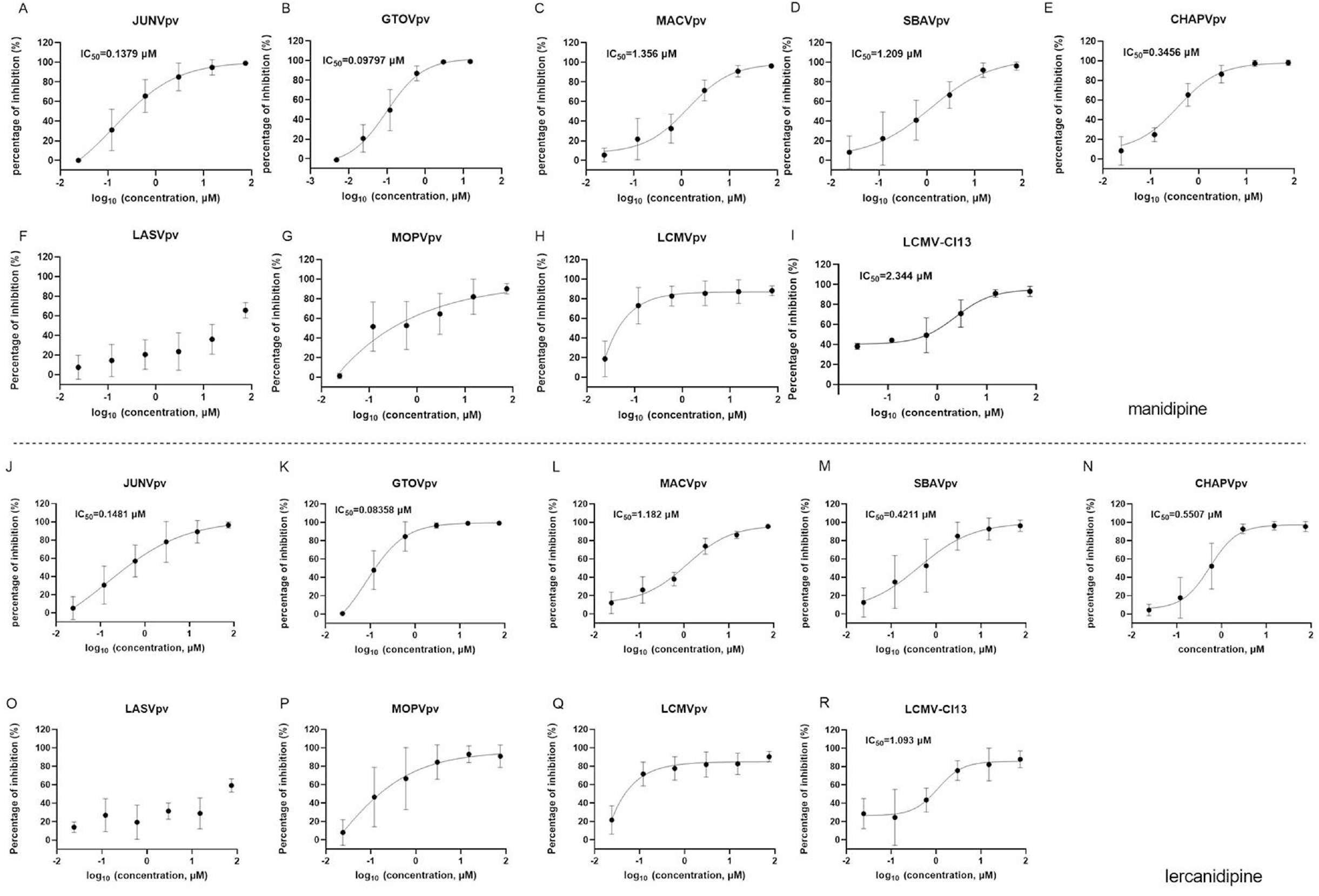
Broad-spectrum antiviral activities of manidipine and lercanidipine. (A-H) Effects of manidipine on the infection of the pseudotype of mammarenaviruses. (I) Inhibition of manidipine against authentic LCMV infection. (J-Q) Effects of lercanidipine on the infection of the pseudotypes of mammarenaviruses. (I) Inhibition of lercanidipine against authentic LCMV infection. Data are presented as means ± SD from three to five independent experiments.

### Manidipine and lercanidipine inhibits LUJV entry by acting as calcium inhibitors

Both manidipine and lercanidipine are DHP VGCC antagonists. VGCC has been reported to be indispensable for the entry of NW mammarenaviruses (20, 21). To this end, we tested the role of small interfering RNAs (siRNAs) in depleting the expression of VGCC genes in U-2 OS cells to determine their effect on LUJV entry; they have been reported to express VGCC genes (*CACNA1S* and *CACNA2D2*) (20, 21). First, siRNA knockdown efficiency was validated using a Western blotting (WB) assay. As shown in Fig. 8A and B, the siRNA knockdown of *CACNA1S*, encoding the VGCC pore-forming α1 subunit targeted by most DHPs. This decreased the protein levels of both CACNA1S and CACNA2D2, whereas the knockdown of *CACNA2DA* decreased CACNA2D2 protein levels. Furthermore, the siRNA knockdown of *CACNA1S* effectively blocked infection by MACVpv, but not by VSVpv; this finding is in line with previous reports that VGCCs are essential for NW mammarenavirus entry (20, 21). Similarly, LUJV infection was inhibited following the knockdown by siRNA of either *CACNA1S* or *CACNA2D2*, indicating that VGCC is critical for LUJV entry (Fig. 8C and D).

**Fig. 8.**
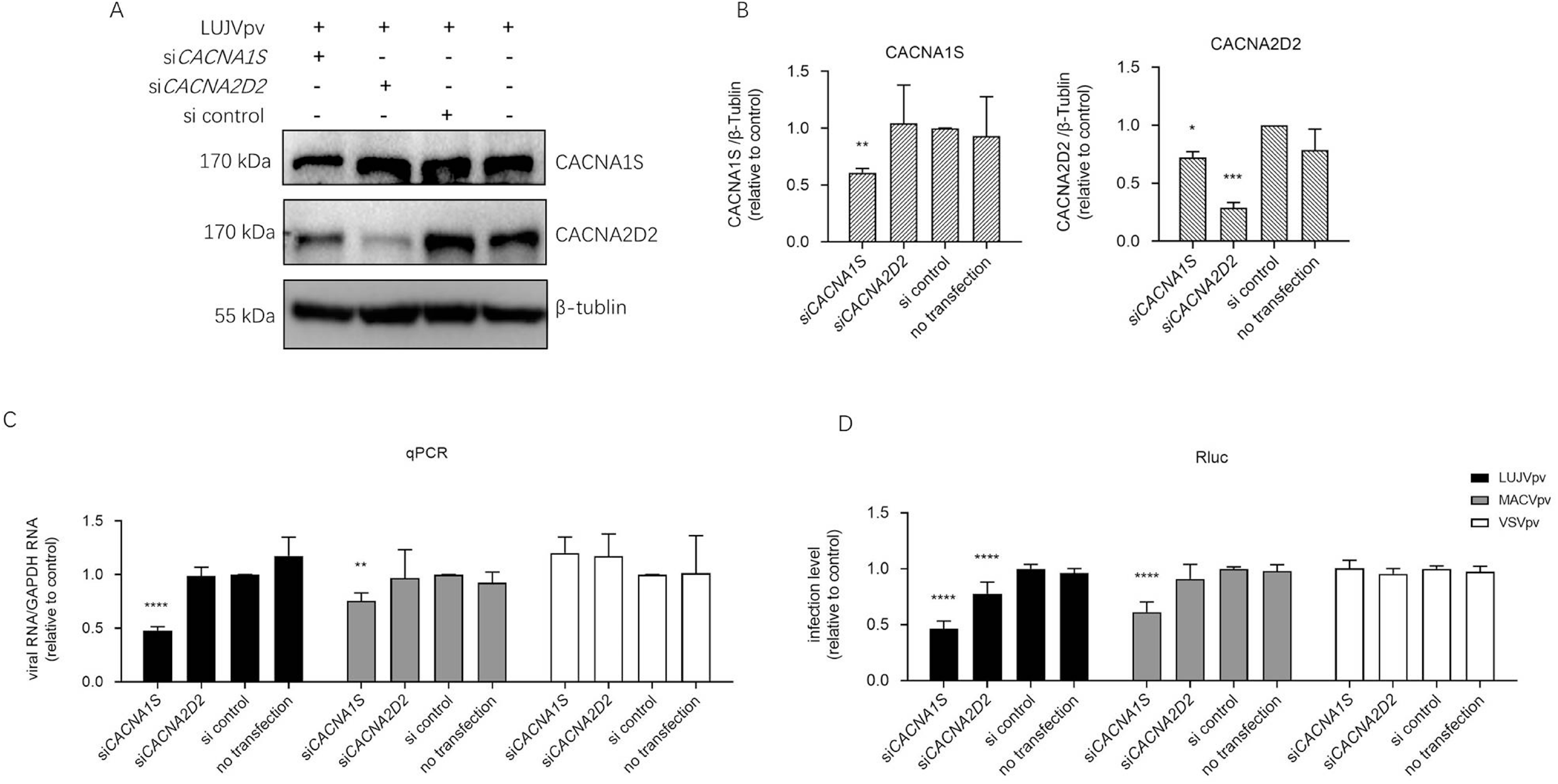
Knockdown of VGCC genes inhibited LUJVpv infection. (A) WB assay of CACNA1S and CACNA2D2 in U-2 OS cells. U-2 OS cells were transfected with si*CACNA1S* and si*CACNA2D2*, respectively. After 48 h, cells were infected with LUJVpv (MOI: 0.1) for 6 h; then, the cell lysates were subjected to a WB assay. (B) Quantification results of WB assay. (C and D) Knockdown of VGCC genes inhibited LUJVpv infection. U-2 OS cells were transfected with si*CACNA1S* or si*CACNA2D2*. After 48 h, cells were infected with LUJVpv (MOI: 0.1), MACVpv (MOI: 0.1), or VSVpv (MOI: 0.1). After 6 h, the duplicate cell lysates were subjected to qPCR (C) and Rluc assays (D), respectively. Data are presented as presented as means ± SD from more than five independent experiments (****, *P* <0.0001; ***, *P* <0.001; **, *P* <0.01; *, *P* <0.05).

## DISCUSSION

Here, we screened an FDA-approved drug library and identified three hit compounds, trametinib, manidipine, and lercanidipine, which prohibited the entry step of pseudotype and recombinant LUJV infections. Among these three hits, trametinib exerted the most effective inhibition, with an IC_50_ of 0.3566 μM and an SI of >560. Trametinib, which is an MAPK inhibitor, is used for to treat patients with unresectable or metastatic melanoma carrying the BRAF V600E mutation. Trametinib has also been shown to play a role as an anti-coronavirus agent that can inhibit the Middle East respiratory syndrome coronavirus infection by modulating the MAPK/ERK signaling pathway (22). It has been shown to inhibit severe acute respiratory syndrome coronavirus 2 (SARS-CoV-2) by down-regulating the expression of ACE2 (23). Intriguingly, investigating the antiviral mechanism of trametinib revealed that it could robustly block LUJV GPC-mediated membrane fusion and its viral target was C410, which was embedded in the TM domain of GP2. In mammarenavirus GPC, the retained SSP interacts with the membrane-proximal external region, as well as the TM domain of GP2. Thus, it stabilizes the pre-fusion conformation of GPC and provides an “Achilles’ heel” that can be targeted by distinct entry inhibitors (13–15, 24–27). To our knowledge, this is the first study to report entry inhibitors targeting the LUJV SSP-GP2 interface, and to identify their viral target(s). Notably, trametinib extended its antiviral spectrum to the pseudotypes of other pathogenic mammarenavirus infections. We demonstrated that trametinib inhibited LASVpv infection by targeting F446 (embedded in the SSP-GP2 interface); it inhibited both the pseudotype and authentic LCMV infection with similar IC_50_ values. For NW mammarenaviruses, we hypothesize that inhibition might occur due to a reduction in the surface expression of the NW mammarenavirus receptor TfR1 (19).

Besides trametinib, the other two hits, manidipine and lercanidipine, were VGCC inhibitors. VGCC inhibitors have been widely reported to inhibit different types of viruses with various antiviral mechanisms (28). Notably, VGCC has been reported to play essential roles in NW mammarenavirus infection (20, 21). This is in line with our results, which showed that both manidipine and lercanidipine robustly inhibited NW mammarenavirus entry, with IC_50_ values ranging from 0.1-1 μM. Lacidipine, which is a VGCC inhibitor, has been demonstrated to inhibit LASVpv and GTOVpv infections, as well as GPC-mediated membrane fusion, by targeting LASV T40 (in the ectodomain of SSP), GTOV V36 (in the ectodomain of SSP), and V436 (in TM domain of GP2) (13). To investigate whether manidipine and lercanidipine inhibit LUJV entry by acting as calcium inhibitors or fusion inhibitors, we studied the effects of both drugs on LUJV GPC-mediated membrane fusion; neither showed promising inhibition of fusion. We tried to select the adaptive mutant by serially passaging with increasing concentrations of both drugs (10–50 μM), but no mutant was observed in GPC. Notably, after reviewing all 29 calcium inhibitors included in the current FDA drug library, we found that besides manidipine and lercanidipine, 12 additional calcium channel blockers (10 μM) inhibited LUJVrv infection to >90% inhibition, suggesting that calcium channels are potential antiviral targets. By using siRNA to knockdown VGCC genes, we found VGCC to be essential for LUJV entry. Based on these results, the calcium channel might serve as a potential antiviral target in LUJV treatment; manidipine and lercanidipine inhibited LUJVpv entry by acting as calcium inhibitors.

The fourth hit compound, ospemifene, is a selective estrogen receptor modulator (SERM); it robustly inhibited LUJV entry in a dose-dependent manner. Notably, during the primary screen all nine SERMs (ospemifene, bazedoxifene HCl, bazedoxifene acetate, clomifene citrate, chlorotrianisene, raloxifene (Evista), tamoxifen, tamoxifen citrate, and toremifene citrate) showed effective inhibition (75-99% inhibition) on LUJVrv infection. However, none of these nine drugs were researched further because of either their cytotoxicity or their inhibition of VSVpv. Notably, a recently published study showed that four SERMs (bazedoxifene HCl, raloxifene (Evista), tamoxifen citrate, and toremifene citrate) could inhibit LUJV infection (29). Similar to our results, these SERMs showed cytotoxicity, with relatively low CC_50_ values (5–10 μM) (29). Although their associated cytotoxicity limits the use of SERMs in antiviral treatment, their promising inhibitory effects suggest that their action mechanisms, that is, the activation or repression of the estrogen target genes (30), might be involved in LUJV infection. Furthermore, the core structures of SERMs might serve as a backbone for structure-activity relationship optimization for drug development.

To date, only one LUJV outbreak has recorded, in 2008; it resulted in a fatality rate of 80% (4/5 cases) (1). There is an urgent need to develop therapeutic options to treat LUJV and other highly pathogenic mammarenavirus infections. Here, we screened an FDA-approved drug library to develop novel therapeutic options for LUJV treatment. Revealing the mechanisms of action of these antiviral inhibitors can facilitate the understanding of LUJV pathogenesis.

## MATERIALS AND METHODS

### Cells and viruses

BHK-21, HEK 293T, Vero, U-2 OS, and A549 cells were cultured in Dulbecco’s modified Eagle’s medium (HyClone, Logan, UT, USA) supplemented with 10% fetal bovine serum (Gibco, Grand Island, NY, USA). The pseudotype VSV, bearing the GPCs of LUJV (GenBank NC_012776.1), LASV (Josiah strain, HQ688673.1), LCMV (Armstrong strain, AY847350.1), MOPV (AY772170.1), GTOV (NC_005077.1), JUNV (XJ13 strain, NC_005081.1), MACV (Carvallo strain, NC_005078.1), SBAV (U41071.1), and CHAPV (NC_010562.1), were generated as previously reported (13, 15). Recombinant VSV expressing the GPCs of LUJV and LASV was generated as described previously (13, 15). The pseudotype and recombinant viruses enveloped by LUJV GPC were designated as LUJVpv and LUJVrv, respectively. The titer of the pseudotype virus was measured by infecting BHK-21 cells that were previously transfected with pCAGGS-VSV G; it was determined by plaque assay 24 h post-infection. The titer of the recombinant virus was determined by a plaque assay. The titers of LUJVrv and LUJVpv were 3.5×10^5^ and 6×10^7^ PFU·ml^−1^, respectively.

### HTS assay of an FDA-approved compound library

A library of 1,775 FDA-approved drugs was purchased from Selleck Chemicals (Houston, TX, USA). The compounds were stored as 10 mM stock solutions in DMSO at −80 °C until use. HTS was performed as shown in Fig. 1A. Vero cells were treated in duplicate with the compounds (10 μM). Then, 1 h later, cells were infected with LUJVrv (MOI, 0.1) for 1 h. After 23 h, cells were fixed with 4% paraformaldehyde and stained with 4’, 6-diamidino-2-phenylindole (DAPI; Sigma-Aldrich, St. Louis, MO, USA). Nine fields per well were imaged on an Operetta high-content imaging system (PerkinElmer) and the percentages of infected and DAPI-positive cells were calculated using the associated Harmony 3.5 software. Cell viability was evaluated using the 3-(4,5-dimethyl-2-thiazolyl)-2,5-diphenyl-2H-tetrazolium bromide (MTT) assay. LUJVpv was used to determine IC_50_ values. Vero cells were treated with compounds of the indicated concentrations. Then, 1 h later, cells were infected with LUJVpv; the supernatant was removed 1 h post-infection. Luciferase activity was measured using the Rluc assay system (Promega, Madison, WI, USA), and the IC_50_ was calculated using GraphPad Prism 8 (GraphPad Inc., La Jolla, CA, USA).

### Membrane fusion assay

293T cells co-transfected with pCAGGS-LUJV GPC, pcDNA 3.1-CD63_GY234AA,_ and pEGFP-N1 were treated with compounds or a vehicle (DMSO) for 1 h, followed by incubation for 20 min in acidified (pH 5.0) medium. The cells were then placed in neutral medium, and syncytium formation was visualized 2 h later via light microscopy. Images were processed using ImageJ software for quantification. The boundaries and coverage areas were traced and calculated using the analyzed particles.

### Selection of the adaptive mutants

Drug-resistant viruses were generated by passaging LUJVrv or LASVrv on vero cells in the presence of 10 μM trametinib. Parallels were passaged in the presence of 2% DMSO as a control. Passaging was terminated when no further improvement in the resistance was detected. RNA from the resistant viruses was extracted using Trizol (TaKaRa, Kusatsu, Shiga, Japan); it was reverse transcribed using the PrimeScript™ RT reagent Kit (TaKaRa). The GPC segment was amplified and sequenced using primers, as previously described (13). Mutant sites were introduced to LUJVpv or LASVpv, as previously described (13), and trametinib sensitivity was determined by Rluc activity.

### RNA interference (RNAi)

To detect the depletion of the target genes in cells, siRNAs from Ambion were used for A1S (siRNA ID: s2297), A2D2 (siRNA ID: 214263), and control (catalog no. 1022076). Briefly, U-2 OS cells were transfected using the forward transfection method of Lipofectamine RNAi Max regent (Invitrogen, Waltham, Massachusetts, USA). SiRNA deletion was carried out for 48 h. Cells were then infected with MACVpv, LUJVpv, and VSVpv (MOI, 0.1) for 6 h. Cell lysates were then subjected to Rluc assay and quantitative PCR (qPCR) assay (primers 5’-GTAACGGACGAATGTCTCATAA-3’ and 5’-TTTGACTCTCGCCTGATTGTAC-3’). All RNA amplifications were normalized to glyceraldehyde 3-phosphate dehydrogenase (GAPDH) RNA (5’-GAAGGTGAAGGTCGGAGTC-3’ and 5’-GAAGATGGTGATGGGATTTC-3’). The antibodies used in the WB assay included anti-CaV1.1 mAB (Invitrogen; 1:1000), anti-CACNA2D2 antibody (Abcam, Cambridge HQ, UK; 1:1000), anti-β-tubulin mouse mAb (AC021; ABclonal, Wuhan, China; 1:1000), horseradish peroxidase (HRP)-linked goat anti-rabbit IgG, and HRP-linked goat anti-mouse IgG ( 1:5000; Proteintech).

## REFERENCES

1. Briese T, Paweska JT, McMullan LK, Hutchison SK, Street C, Palacios G, Khristova ML, Weyer J, Swanepoel R, Egholm M, Nichol ST, Lipkin WI. 2009. Genetic detection and characterization of Lujo virus, a new hemorrhagic fever-associated arenavirus from southern Africa. PLoS Pathog 5:e1000455.

2. Oldstone MB. 2002. Arenaviruses. I. The epidemiology molecular and cell biology of arenaviruses. Introduction. Curr Top Microbiol Immunol 262:V–XII.

3. Nunberg JH, York J. 2012. The curious case of arenavirus entry, and its inhibition. Viruses 4:83–101.

4. Cohen-Dvashi H, Kilimnik I, Diskin R. 2018. Structural basis for receptor recognition by Lujo virus. Nat Microbiol 3:1153–1160.

5. Raaben M, Jae LT, Herbert AS, Kuehne AI, Stubbs SH, Chou YY, Blomen VA, Kirchhausen T, Dye JM, Brummelkamp TR, Whelan SP. 2017. NRP2 and CD63 Are Host Factors for Lujo Virus Cell Entry. Cell Host Microbe 22:688–696 e5.

6. Buchmeier MJ, de la Torre JC, Peters CJ. 2007. Fields Virology, 5th ed. Lippincott-Raven, Philadelphia.

7. Vela E. 2012. Animal models, prophylaxis, and therapeutics for arenavirus infections. Viruses 4:1802–29.

8. Eichler R, Lenz O, Strecker T, Garten W. 2003. Signal peptide of Lassa virus glycoprotein GP-C exhibits an unusual length. FEBS Lett 538:203–6.

9. Wang W, Zhou Z, Zhang L, Wang S, Xiao G. 2016. Structure-function relationship of the mammarenavirus envelope glycoprotein. Virol Sin 31:380–394.

10. Igonet S, Vaney MC, Vonrhein C, Bricogne G, Stura EA, Hengartner H, Eschli B, Rey FA. 2011. X-ray structure of the arenavirus glycoprotein GP2 in its postfusion hairpin conformation. Proc Natl Acad Sci U S A 108:19967–72.

11. Lenz O, ter Meulen J, Klenk HD, Seidah NG, Garten W. 2001. The Lassa virus glycoprotein precursor GP-C is proteolytically processed by subtilase SKI-1/S1P. Proc Natl Acad Sci U S A 98:12701–5.

12. Garbutt M, Liebscher R, Wahl-Jensen V, Jones S, Moller P, Wagner R, Volchkov V, Klenk HD, Feldmann H, Stroher U. 2004. Properties of replication-competent vesicular stomatitis virus vectors expressing glycoproteins of filoviruses and arenaviruses. J Virol 78:5458–65.

13. Wang P, Liu Y, Zhang G, Wang S, Guo J, Cao J, Jia X, Zhang L, Xiao G, Wang W. 2018. Screening and Identification of Lassa Virus Entry Inhibitors from an FDA-Approved Drugs Library. J Virol 92:e00954–18.

14. Zhang G, Cao J, Cai Y, Liu Y, Li Y, Wang P, Guo J, Jia X, Zhang M, Xiao G, Guo Y, Wang W. 2019. Structure-activity relationship optimization for lassa virus fusion inhibitors targeting the transmembrane domain of GP2. Protein Cell 10:137–142.

15. Liu Y, Guo J, Cao J, Zhang G, Jia X, Wang P, Xiao G, Wang W. 2021. Screening of Botanical Drugs against Lassa Virus Entry. J Virol doi:10.1128/JVI.02429-20.

16. Cao J, Zhang G, Zhou M, Liu Y, Xiao G, Wang W. 2021. Characterizing the Lassa Virus Envelope Glycoprotein Membrane Proximal External Region for Its Role in Fusogenicity. Virol Sin 36:273–280.

17. Veit M. 2012. Palmitoylation of virus proteins. Biol Cell 104:493–515.

18. Ito H, Watanabe S, Takada A, Kawaoka Y. 2001. Ebola virus glycoprotein: proteolytic processing, acylation, cell tropism, and detection of neutralizing antibodies. J Virol 75:1576–80.

19. Horniblow RD, Bedford M, Hollingworth R, Evans S, Sutton E, Lal N, Beggs A, Iqbal TH, Tselepis C. 2017. BRAF mutations are associated with increased iron regulatory protein-2 expression in colorectal tumorigenesis. Cancer Sci 108:1135–1143.

20. Sarute N, Ross SR. 2020. CACNA1S haploinsufficiency confers resistance to New World arenavirus infection. Proc Natl Acad Sci U S A doi:10.1073/pnas.1920551117.

21. Lavanya M, Cuevas CD, Thomas M, Cherry S, Ross SR. 2013. siRNA screen for genes that affect Junin virus entry uncovers voltage-gated calcium channels as a therapeutic target. Sci Transl Med 5:204ra131.

22. Kindrachuk J, Ork B, Hart BJ, Mazur S, Holbrook MR, Frieman MB, Traynor D, Johnson RF, Dyall J, Kuhn JH, Olinger GG, Hensley LE, Jahrling PB. 2015. Antiviral potential of ERK/MAPK and PI3K/AKT/mTOR signaling modulation for Middle East respiratory syndrome coronavirus infection as identified by temporal kinome analysis. Antimicrob Agents Chemother 59:1088–99.

23. Zhou L, Huntington K, Zhang S, Carlsen L, So EY, Parker C, Sahin I, Safran H, Kamle S, Lee CM, Geun Lee C, J AE, K SC, M TN, W JA, Youssef E, J AP, Navaraj A, A AS, Liang O, El-Deiry WS. 2020. MEK inhibitors reduce cellular expression of ACE2, pERK, pRb while stimulating NK-mediated cytotoxicity and attenuating inflammatory cytokines relevant to SARS-CoV-2 infection. Oncotarget 11:4201–4223.

24. Shankar S, Whitby LR, Casquilho-Gray HE, York J, Boger DL, Nunberg JH. 2016. Small-Molecule Fusion Inhibitors Bind the pH-Sensing Stable Signal Peptide-GP2 Subunit Interface of the Lassa Virus Envelope Glycoprotein. J Virol 90:6799–807.

25. Tang K, Zhang X, Guo Y. 2020. Identification of the dietary supplement capsaicin as an inhibitor of Lassa virus entry. Acta Pharm Sin B 10:789–798.

26. Lee AM, Rojek JM, Spiropoulou CF, Gundersen AT, Jin W, Shaginian A, York J, Nunberg JH, Boger DL, Oldstone MB, Kunz S. 2008. Unique small molecule entry inhibitors of hemorrhagic fever arenaviruses. J Biol Chem 283:18734–42.

27. Larson RA, Dai D, Hosack VT, Tan Y, Bolken TC, Hruby DE, Amberg SM. 2008. Identification of a broad-spectrum arenavirus entry inhibitor. J Virol 82:10768–75.

28. Chen X, Cao R, Zhong W. 2019. Host Calcium Channels and Pumps in Viral Infections. Cells 9.

29. Welch SR, Spengler JR, Genzer SC, Chatterjee P, Flint M, Bergeron E, Montgomery JM, Nichol ST, Albarino CG, Spiropoulou CF. 2021. Screening and Identification of Lujo Virus Inhibitors Using a Recombinant Reporter Virus Platform. Viruses 13.

30. Riggs BL, Hartmann LC. 2003. Selective estrogen-receptor modulators -- mechanisms of action and application to clinical practice. N Engl J Med 348:618–29.

